# Role of the inner membrane cytochrome ImcH in *Geobacter* extracellular electron transfer and energy conservation

**DOI:** 10.1101/2023.05.03.539133

**Authors:** Andreia I. Pimenta, Catarina M. Paquete, Leonor Morgado, Marcus J. Edwards, Thomas Clarke, Carlos A. Salgueiro, Inês A.C. Pereira, Américo G. Duarte

## Abstract

Electroactive bacteria combine the oxidation of carbon substrates with an extracellular electron transfer (EET) process that discharges electrons to an electron acceptor outside the cell. This process involves electron transfer through consecutive redox proteins that efficiently connect the inner membrane to the cell exterior. In this study, we isolated and characterized the quinone-interacting membrane cytochrome *c* ImcH from *Geobacter sulfurreducens*, which is involved in the EET process to high redox potential acceptors. Our work provides evidence that ImcH is electroneutral, as it transfers electrons and protons to the same side of the membrane, contributing to the maintenance of a proton motive force, and plays a central role in recycling the menaquinone pool.

**Importance:** *Geobacter sulfurreducens* is a model electroactive bacterium, widespread in the environment and of significant interest for biotechnological applications. Its ability to form thick and conductive biofilms on top of conducting surfaces makes this microbe very useful in bioelectrochemical systems for the production of energy or added value products. To explore *Geobacter* spp. as a biocatalyst it is essential to understand its metabolism, particularly the molecular mechanisms for extracellular electron transfer and energy conservation. Our results reveal the importance of ImcH in both processes, identifying this protein as a major player on *Geobacter* metabolism.

## Introduction

Microbial communities can live in almost any environment, and the permanence of each microorganism relies on its metabolic ability to use organic or inorganic molecules that are present in the environment, which can be inside or outside the cell, in order to produce energy. Electroactive microorganisms can exchange energy with the environment and with other microorganisms through a process called extracellular electron transfer (EET). In this process, the electrons released from catabolic reactions are transported to outside of the cell to a terminal acceptor, which can be a mineral, an electrode or another microorganism in a process named direct interspecies electron transfer (DIET) [1]. These microorganisms have been the focus of research for many years, with the goal of understanding and optimizing EET processes for biotechnological applications, including bioremediation, bioelectronics and bioelectrochemical devices [2,3]. *Geobacter sulfurreducens* is frequently found in bioelectrochemical systems, in mixed or pure cultures, due to its ability of assemble thick and conductive biofilms, either to discharge electrons to an electrode surface or receive electrons for bioelectrosynthesis. Formation of such conductive biofilms is a main advantage of *Geobacter*, which involves the electron transfer over micrometers by assembling unique oligomeric structures of cytochrome nanowires, such as OmcS [4], OmcZ [5] and OmcE [6].

*G. sulfurreducens* is metabolically versatile, and can couple the oxidation of acetate, formate and hydrogen (H_2_) to the reduction of soluble and insoluble minerals [7,8], or electrodes at a wide range of reduction potentials, and produce energy through chemiosmotic ATP synthesis [9]. The oxidation of carbon substrates or H_2_ and electron transfer to the menaquinone (MK) pool is the starting point for EET and energy conservation, which allows the establishment of a proton motive force (*pmf*), required for ATP synthesis. To oxidize the menaquinol (MKH_2_)-pool, *G. sulfurreducens* has several menaquinol oxidorreductases [10], but only three were proven to be essential for EET, ImcH, CbcL and CbcBA. ImcH is a multiheme cytochrome *c* with an N-terminal membrane domain with three transmembrane helices (TMH) and a C-terminal domain with seven *c*-type heme binding motifs, which was demonstrated to be essential for EET to high potential electron acceptors such as soluble Fe citrate, or electrodes poised at potentials above -100 mV [11]. CbcL is an inner membrane protein that has an N-terminal soluble domain with nine heme *c* binding motifs and a C-terminal membrane domain homologous to di-heme cytochrome *b* subunits. Contrary to ImcH, CbcL was proven to be essential for EET to insoluble Fe oxides with low redox potentials, or electrodes poised at -250 mV [12,13]. CbcBA is an inner membrane complex where CbcA is a multiheme cytochrome *c* subunit predicted to carry seven *c*-type hemes in the N-terminal domain with a single TMH in the C-terminal domain, and CbcB, an integral membrane subunit homologous to the di-heme cytochrome *b* subunits with four TMH. This complex is important for EET towards minerals with a lower potential and was proven to be essential in conditions close to the thermodynamic limit of respiration when acetate is the electron donor [14]. To link the inner and outer membranes, *Geobacter* species express a high level of cytochromes *c* in the periplasm, namely the Ppc family of triheme cytochromes with five homologues, of which PpcA is the most abundant in the *Geobacter* periplasm, and its deletion from the genome strongly impairs Fe(III) respiration [15]. At the outer membrane several protein complexes composed by an integral β-barrel pore, a periplasmic and an outer membrane multiheme cytochrome such as the OmaB-OmbB-OmcB, ExtABCD, or their homologues allow for an efficient EET towards the outside [16]. It has been proposed that the excess of cytochromes in the periplasm may function as a capacitor, storing electrons from the metabolism, as their lack of specificity enables fast electron transport from the inner towards the outer membrane [17].

In this work, we used a combination of biochemical, spectroscopic, electrochemical and computational methods to understand the role of the MK-interacting ImcH in *G. sulfurreducens* energy metabolism and in the EET process.

## Results

### ImcH biochemical and spectroscopic characterization indicates it is a dimer in solution

Recombinant ImcH was successfully produced in *Shewanella oneidensis* and purified (Figure 1A) as a protein with a molecular mass of ∼66 kDa, which is in agreement with the predicted MW = 65 kDa (recombinant protein with seven hemes). N-terminal sequencing (MTLRKT) confirmed that no signal peptide was cleaved, as predicted by bioinformatic tools, while heme iron quantification determined a 6.6 iron atoms to protein ratio, consistent with the presence of seven *c*-type hemes as predicted from the number heme *c* binding motifs in the sequence. Size exclusion chromatography of recombinant ImcH solubilized in *n*-dodecyl-β-D-maltoside (DDM) shows a molecular mass of 121 (± 27) kDa, consistent with a dimeric form of the protein. On the other hand, sedimentation velocity c(S) distribution analysis through repeated measurement of the absorbance (410 nm) during centrifugation shows a heterogenous distribution with two major peaks, which correspond to molecular masses of 165 (± 18) kDa and 312 (± 40) kDa (Figure 1B) with equal proportion, consistent with the presence of a dimer and a tetramer, respectively. The same experiments performed with ImcH solubilized in Triton X-100 shifted the c(S) distribution to lower values with molecular masses of 112 (± 7) kDa and 215 (± 20) kDa (supplementary information figure S1). Triton X-100 has a partial specific volume (v-bar) of 0.91 ml g^−1^ so at a solvent density of 1.099 g ml^−1^ Triton will be effectively weightless and not contribute to the ImcH buoyant mass. c(S) analysis of ImcH in buffer containing Triton X-100 and 0-60 % D_2_O caused a linear decrease in the observed sedimentation coefficient (supplementary information figure S1). Extrapolation allows the observed sedimentation coefficient to be predicted at the density that matches the buoyancy of Triton X-100. Assuming the ImcH protein is a globular protein, the predicted sedimentation coefficient of 3.43 S would correspond to a protein of 68 kDa. This demonstrates that the detergent used for ImcH solubilization clearly influences the oligomerization state of the protein between a monomer and a dimer.

**Figure 1.**
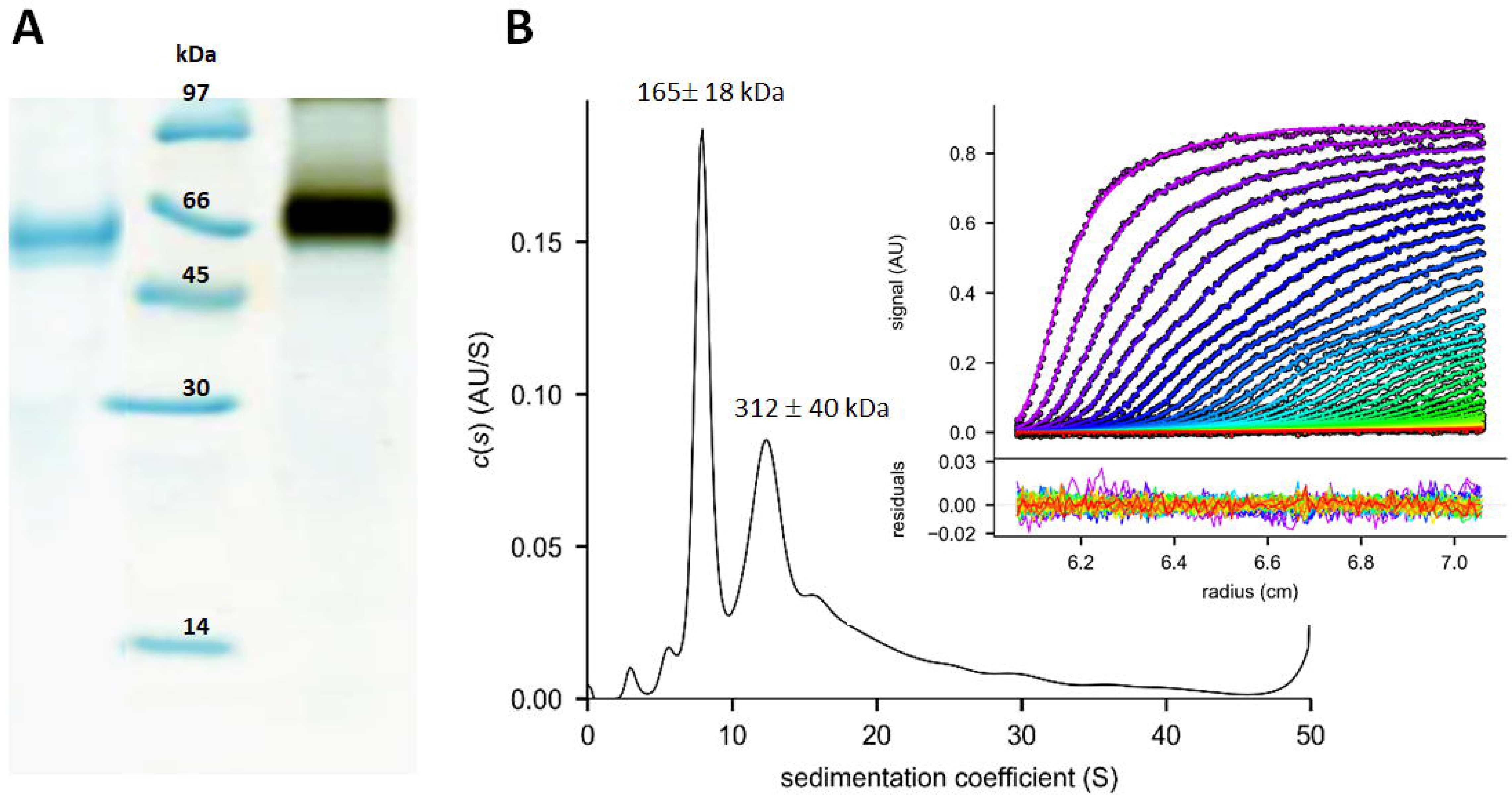
– *G. sulfurreducens* ImcH biochemical characterization. (A) SDS-Page electrophoresis of pure ImcH stained for total protein (left) and heme staining (right), Protein marker - low molecular weight standards (Cytiva). (B) Sedimentation velocity analysis of ImcH, inset show the monitored absorbance at 410 nm (markers) and fit (lines) to the Lamm equation with residual absorption from the fitted data (lower inset panel) The fitted c(s) distribution of sedimentation coefficients indicates the presence of two species.

### ImcH has an unusual predicted His/Gln heme coordination

An ImcH structural model was constructed using AlphaFold2 and the *c*-type hemes were manually fitted to the cysteine residues of the corresponding CXXCH heme binding motifs, with the histidine as the proximal heme Fe ligand (Figure 2). The ImcH model displays a membrane domain with three TMH at the N-terminal (composed by 130 aminoacids), followed by a soluble domain (aminoacids 131 to 507) that accommodates the seven heme co-factors (Figure 2A). The model shows that hemes 1 and 7 (numbering according to the position of the heme binding motif at the ImcH primary sequence) present higher solvent exposure, with the first near the hydrophobic domain and the last on the opposite side of the soluble domain of the structure. The seven hemes are in close proximity with Fe-to-Fe distances lower than 12 Å, and disposed in an inverted L-shape (Figure 2B) with different dihedral angles between the ligands. All redox centers show *bis*-His coordination apart from the first heme, which seems to be coordinated by a histidine (H137) and has a glutamine (Q179) in the second coordination position (Figure 2C). A multiple sequence alignment reveals that the heme binding motifs and residues for the Fe coordination (including Q179) are conserved among ImcH homologues (supplementary information Figure S2).

**Figure 2.**
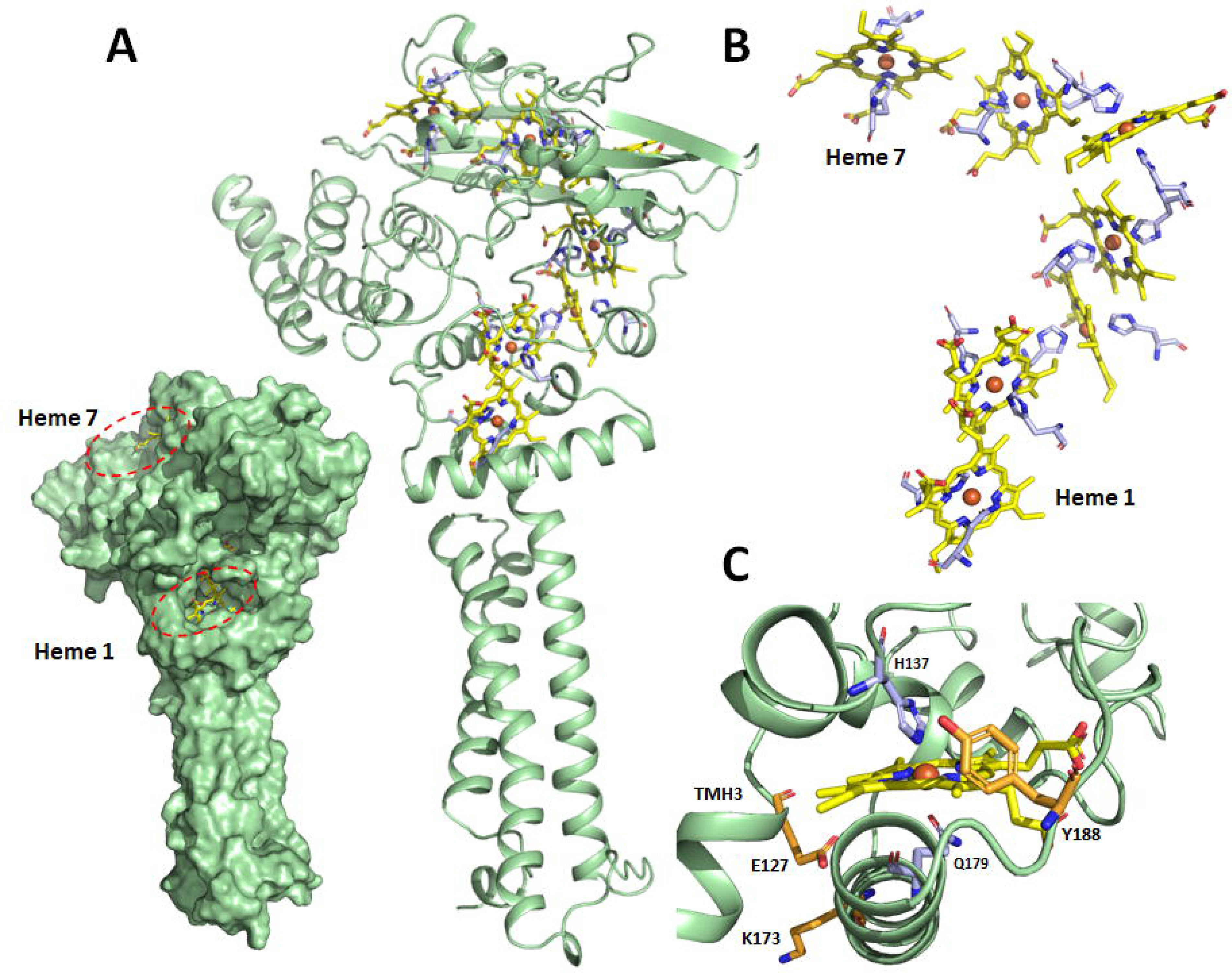
– *G. sulfurreducens* ImcH model. (A) The structural model of ImcH as surface (left) or as a cartoon (right). The hemes are represented as sticks with carbons colored in yellow, nitrogen in blue, oxygen in red, and Fe atoms as orange spheres). (B) ImcH heme arrangement with its correspondent Fe axial ligands represented as sticks (carbons colored in light blue, nitrogen in blue, oxygen in red). (C) ImcH heme 1 showing the unusual His/Gln coordination and conserved protonable residues represented as sticks (carbons colored in light orange, nitrogen in blue, oxygen in red).

ImcH is homologous to members of the NapC/NirT/NrfH family despite the different number of TMH and redox co-factors [11],. A structural superposition of the ImcH model and the NrfH structure, the membrane anchor subunit of the menaquinol-interacting cytochrome *c* nitrate reductase (NrfA_4_H_2_), which is the only structure available within the NapC/NirT family [18], shows a similar structure between the two proteins, with a close overlap of one of the TMH and the first four ImcH hemes (supplementary figure S3). Similarly to ImcH, heme 1 from NrfH contains a unique coordination: a methionine from the sequence CXXCHXM is replacing the canonical histidine residue that acts as the proximal axial ligand, and on the other side of the heme an aspartate residue is at coordinating distance to heme-Fe, but no electron density for a bond was observed [18]. In NrfH, the menaquinone analogue 2-heptyl-4-quinolinol 1-oxide (HQNO)was shown to binds to a pocket close to this heme pocket, in direct contact with the lipid layer [18]. In ImcH model, an identical structural arrangement is observed, where heme 1 is shielded from the solvent by a helix that lays almost perpendicular to the transmembrane domain (Figure 2 and supplementary figure S3). Based on the ImcH model, it seems that these two proteins have similar menaquinone-binding sites, close to heme 1, immediately following the transmembrane domain, and with an unusual coordination. In NrfH, a single TMH is positioned on the back of the quinone molecule, and a similar organization is observed in ImcH, where TMH3 overlaps with this NrfH TMH. ImcH E127 from this helix, and K173 and Y188, which are in the vicinity of this heme pocket (Figure 2C), are highly conserved also among ImcH homologues (supplementary figure S2), suggesting that they can be involved in MKH_2_ oxidation, possibly in proton transport to the P-side of the membrane. In *Wolinella succinogenes* NrfH replacement of the K83 by a non protonable residue (isoleucine) prevents the cells from reducing nitrite to ammonia [19]. This lysine (Lys 82, in *D. vulgaris)* overlaps with K173 in the ImcH model supplementary Figure S3.

### ImcH spectroscopic features

UV-visible spectra of purified ImcH (Figure 3A) display a typical cytochrome signature with an intense Soret band centered at 410 nm that shifts to 421 nm upon reduction, with an additional increase of absorbance in the characteristic α and β bands centered at 523 and 552 nm, respectively. The EPR spectrum (Figure 3B) shows a complex set of resonances characteristic of low-spin (S=1/2) *bis*-His coordinated hemes. These centers can present different dihedral angles between the imidazole coordinating ligands and possible spin-spin interactions due to a close proximity of the centers, both predicted by the ImcH model, and observed in other multiheme cytochromes [20–22]. The resonance at g value of 5.99 indicates the presence of a partial high-spin heme (S=5/2). This signal may be attributed to ImcH heme 1, as observed for homologous proteins [23], where the unusual His/Gln coordination may not be sufficiently stable to ensure an homogeneous low-spin coordination. Interestingly, a glutamine heme coordination was described in single mutant variants of the yeast cytochrome *c* peroxidase H175Q [24] and in *Bacillus megaterium* P450 flavocytochrome variant A264Q, where in this last case, a Cys/Gln heme-Fe showed a rhombic low-spin EPR signal, but the Gln coordination was easily displaced by the presence of NO [25]. The low intensity of the high-spin signal compared to the low-spin signatures may arise from possible magnetic coupling of the redox centers, which was observed in several multiheme cytochromes [22,23,26], and may also occur in ImcH due to its dimerization when solubilized in DDM. The spin-state of ImcH in DDM was also evaluated by 1D ^1^H NMR in the oxidized and reduced forms (Supplementary figure S4). However, no signal was observed in the regions typical for heme resonances in either redox states probably due to the high molecular weight of the protein dimer solubilized in detergent.

**Figure 3.**
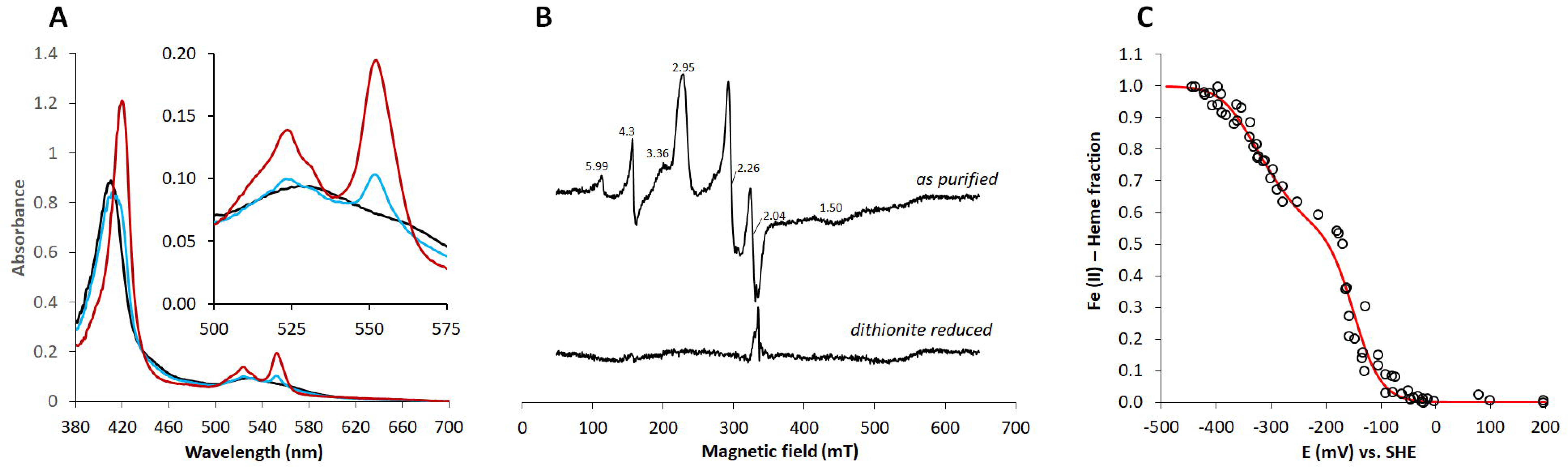
– ImcH spectroscopic characterization. (A) UV-visible spectra of pure ImcH, black trace-as purified (oxidized), blue trace-reduced with excess of menadiol and red trace-reduced with excess of sodium dithionite. (B) EPR spectra of the ImcH: upper - as purified (oxidized), and lower - reduced with excess of sodium dithionite. (C) UV-visible potentiometric titration of ImcH, open circules - experimental data, red curve - theoretical fit for the sum of seven n=1 Fe^2+^/Fe^3+^ redox centers.

### ImcH is redox active between -350 and -150 mV

To determine the redox properties of ImcH, menadiol, a soluble menaquinone analogue, was used to reduce DDM-solubilized ImcH. UV-visible spectroscopy showed that an excess of menadiol was not enough to fully reduce ImcH (Figure 3A), since reduction was only achieved for approximately two of the seven hemes present in the protein. This incomplete reduction has been previously reported for CymA, another member of the NapC/NirT/NrfH family, essential for menaquinol oxidation in *Shewanella* species [23]. To investigate the range of potentials where ImcH is redox active, an UV-visible potentiometric titration was performed for the reductive and oxidative directions (Figure 3C). The experimental data simulated with the sum of seven Fe^2+^/Fe^3+^ (n=1) redox centers predicts apparent midpoint reduction potentials of four electron transfer processes at -150 mV (vs SHE), and single events at -274, -326 and -358 mV (vs SHE). These reduction potentials cannot be assigned to specific redox centers in the structure, since this methodology does not allow to discriminate the individual hemes [27]. However these values are in agreement with the incomplete reduction of ImcH with excess of menadiol (E^0^=-70 mV), and its complete reduction in the presence of dithionite (E^0^ ≅ -470 mV) [28], as observed by EPR spectroscopy.

### ImcH interacts with the Ppc triheme cytochrome family

ImcH faces the periplasm of *G. sulfurreducens* where several soluble cytochromes can act as possible physiological partners. It has been demonstrated that the triheme Ppc cytochrome family, composed by five homologues (PpcA, PpcB, PpcC, PpcD and PpcE) are involved in periplasmic electron transfer in *Geobacter* and are physiologically equivalent [17]. These proteins show similar structure but differ in expression conditions [17] and heme reduction potentials [29–31]. Besides these triheme cytochromes, *G. sulfurreducens* expresses a periplasmic monohemic cytochrome PccH that was proposed to be involved in EET processes, and is essential for the reverse electron flow in current consuming biofilms [32]. To investigate the possible interacting partners of ImcH, Surface Plasmon Resonance (SPR) experiments were performed with recombinant ImcH immobilized in a nitrilotriacetic acid sensor chip through its C-terminal 6x His-tag, and the interaction with the cytochromes PpcA, PpcB, PpcD, PpcE and PccH were evaluated (Figure 4A). All the cytochromes tested interact with ImcH, with PpcA showing the highest response. Although steady-state conditions could not be achieved, it was possible to estimate an equilibrium dissociation constant (K^D^) of 2.7 ± 0.5 μM, based on the maximum response derived from duplicate sensorgrams (Figure 4B).

**Figure 4.**
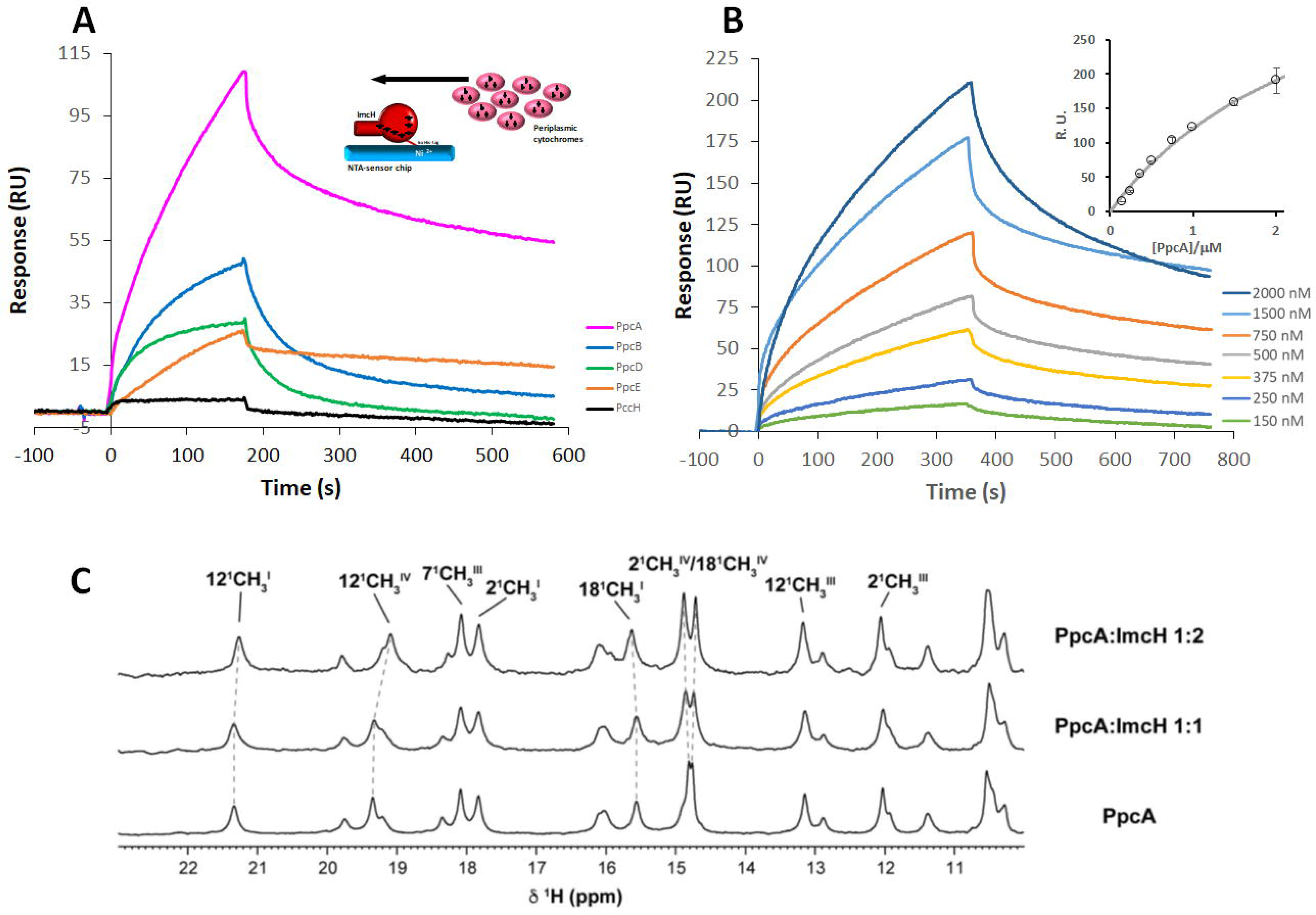
– Interaction between ImcH and PpcA. (A) Sensorgrams of ImcH interaction with PpcA, PpcB, PpcD, PpcE and PccH. (B) Sensorgrams of ImcH interaction with increasing concentrations of PpcA (from 150 to 2000 nM); Inset presents the maximum steady state response (average of two replicates) as function of PpcA concentration to determine the dissociation constant. (C) Low-field 1D ^1^H NMR spectra for the interaction between ImcH and PpcA, with increasing molar ratio. The dashed lines follow the chemical shift changes of the resonances that show higher variation in the presence of ImcH, which are mapped on PpcA structure (PDB 2MZ9) in supplementary figure S5.

1D ^1^H NMR experiments were performed to obtain further insights on the interaction between ImcH and PpcA (Figure 4C). The mixture of PpcA and ImcH in equimolar (1:1), or in twice-molar excess of ImcH (1:2) lead to perturbations on the PpcA spectrum, specifically on heme I and IV methyl groupś resonances (Supplementary Figure S5), suggesting a binding site between these two hemes. Together, SPR and NMR data suggest that the abundant PpcA is most likely the physiological partner of ImcH.

### ImcH transfers electrons to PpcA

Electron transfer between ImcH and PpcA was further studied by cyclic voltammetry (CV) and 1D ^1^H NMR experiments. For the CV experiments, ImcH was immobilized to a stationary pyrolytic graphite electrode and the electron transfer to PpcA was evaluated upon its addition to the electrolyte solution. Cyclic voltammogram subtraction of experiments done with and without PpcA in solution show the development of a cathodic peak centered at -170 mV that derives from the continuous electron transfer from the electrode-bound ImcH to PpcA (Figure 5A). Control experiments with the bare electrode with and without an equivalent concentration of PpcA in solution show a lower current signal with an asymmetric quasi-reversible redox process that has an apparent midpoint redox potential of -85 mV (Figure 5A). This process is attributed to the redox interaction of PpcA at the electrode surface, and is in between PpcA redox potentials (-171 and - 60 mV, at pH 7.0 – see supplementary table S1). Cyclic voltammograms that confirm ImcH adsorption to the electrode, and the ones used for subtraction are provided in supplementary Figure S6.

**Figure 5.**
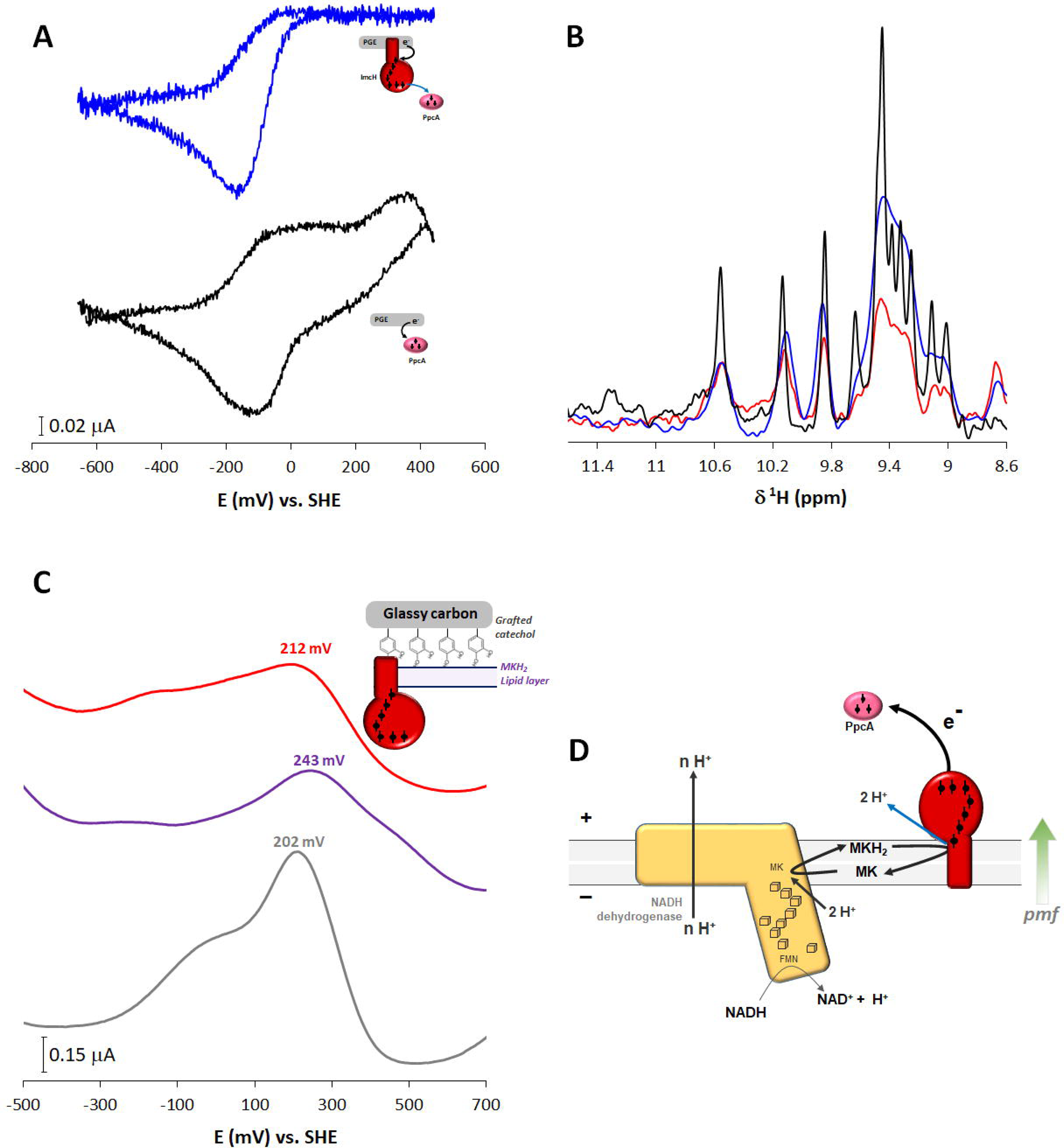
– Direct electron transfer between ImcH and PpcA. (A) Cyclic voltammogram (100 mV s^−1^) subtraction of ImcH modified graphite electrode with 1.5 µM of PpcA in the electrolyte solution minus the cyclic voltammogram obtained without PpcA in the electrolyte solution (blue) bare graphite electrode with 1.5 µM of PpcA in the electrolyte solution minus the cyclic voltammogram obtained without PpcA in the electrolyte solution (black). The cyclic voltammograms used for subtraction are provided in supplementary figure S6. (B) Low-field 1D ^1^H NMR spectra to detect the interaction between reduced ImcH and oxidized PpcA at increasing molar ratios, fully reduced PpcA (black), mixture of reduced ImcH with oxidized PpcA at 1:1 (red) and 1:2 molar ratio (blue). Individual spectra are provided in supplementary figure S7. (C) Differential pulsed voltammograms for the catechol grafted glassy carbon electrode bare electrode (grey), electrode coated with a MKH_2_/lipids mixture (purple) and electrode coated with the same mixture with ImcH solubilized (red). (D) Schematic representation of the redox loop mechanism involved in *G. sulfurreducens* energy conservation for reduction of high potential electron acceptors. NADH dehydrogenase (Complex I) recycles NADH produced from acetate oxidation and drives electrons to ImcH for EET; FMN - flavin mononucleotide co-factor; - cubane Fe-S center; - heme co-factors.

In addition, the electron transfer reaction between ImcH and PpcA was also studied by 1D ^1^H NMR titration experiments. PpcA has a characteristic and unique 1D ^1^H NMR spectra in each oxidation state (supplementary figure S7). Due to the paramagnetic effect of the heme irons, the heme resonances spread in a larger spectral range in the oxidized state in contrast to the reduced state [33]. Also, due to the presence of several heme groups, these signals at intermediate oxidation states are broader and are found in chemical shifts ranges between the fully reduced and fully oxidized states. Consequently, it is possible to monitor any change in the oxidation state of PpcA upon its addition to reduced ImcH under anaerobic conditions (Figure 5B) [33]. Following the addition of oxidized PpcA to reduced ImcH, signals that are characteristic of the reduced PpcA appear, confirming the electron transfer between the two proteins. The addition of increasing amounts of oxidized PpcA led to broadening of this set of signals (1:2 molar ratio) and to the appearance of broad and weak resonances at lower field ∼ 12 ppm (1:3 molar ratio), which are characteristic of PpcA intermediate oxidation states [29,34] (supplementary figure S7). At these protein ratios both ImcH and PpcA remain in intermediate oxidation states, and the results provide evidence for electron equilibration between the two proteins, as expected from the apparent macroscopic redox potentials of -358 to -150 mV for ImcH and -171 to -60 mV for PpcA (see supplementary Table S1).

### ImcH transfers protons from MKH_2_ to the periplasm

ImcH is described as a MKH_2_:oxidoreductase that oxidizes the MKH_2_ pool and transfers electrons to the periplasm for further EET. So far, no information is available regarding the direction of proton release upon MKH_2_ oxidation. To study this, differential pulsed voltammetry experiments were performed using a catechol grafted glassy carbon electrode. In this setup, catechol works as a pH sensitive probe, since its midpoint redox potential decreases with an increase of pH [35]. The catechol grafted bare electrode shows a peak centered at 202 mV that shifts to 243 mV when the electrode is modified with a mixture of lipids containing MKH_2_ (Figure 5C). This change is due to the acidification of the electrode vicinity with the protons released from MKH_2_ oxidation by the electrode. Incorporation of ImcH on top of this MKH_2_-lipid layer shows that the potential measured (212 mV) changes to a value that is closer to the catechol redox potential when grafted to the bare electrode (202 mV). Considering that ImcH likely orients at the electrode-deposited bilayer with its hydrophilic domain facing the solution, this result indicates that protons released from MKH_2_ oxidation by ImcH are transfered away from the electrode, and therefore the grafted catechol does not detect their presence. As observed in the ImcH model, the solvent exposed heme 1 pocket that is the likely MKH_2_-binding site, contains conserved protonatable residues that face the P-side of the membrane and can be involved in the transfer of protons from the quinone binding site to the periplasm. These results classify ImcH as electroneutral, since it releases electrons and protons to the same side of the membrane.

## Discussion

The quinone-interacting ImcH is widespread in electroactive bacteria from different classes, such as Acidobacteria, Bacteroidota, Ignavibacteriae, Verrucomicrobia and Deltaproteobacteria [36]. In *Geobacter* species all the identified ImcH homologues have three predicted TMH and seven heme binding motifs, while in other bacteria this may differ, with homologues with only one or two TMH and with five to nine hemes [36]. ImcH structural features resemble members of the NapC/NirT family, including the MK-interacting anchor subunits of the nitrite reductase (NrfH), nitrate reductase (NapC), DMSO reductase (DorC), TMAO reductase (TorC) and the inner membrane cytochrome CymA, which is present in Beta and Gammaproteobacteria metal reducers and oxidizers [37]. NrfH, NapC, DorC and TorC transfer electrons from the MKH_2_ pool to a physiologic enzymatic partner in the periplasm (i.e. NrfA, NapAB, DorA and TorA, respectively), with which it may assemble in a stable complex. CymA, on the other hand, is less specific concerning redox partners and it can transfer electrons to several redox proteins in the periplasm of *Shewanella*, allowing this organism to respire distinct soluble and insoluble compounds [38–41].

The transfer of electrons outside the cell poses problems that are not present when the terminal electron acceptor is inside the cell, namely it requires a continuous electron-transfer pathway to connect the inner membrane to the exterior. Our work demonstrates that ImcH can directly reduce periplasmic Ppc cytochromes, and protein-protein interaction experiments show a higher affinity towards PpcA over its homologues. No interaction was detected with the monohemic PccH that has a much higher redox potential, which could favor the electron transfer process. CbcL, which is required for the respiration of terminal acceptors with a low redox potential also recruits PpcA for EET [42]. This does not exclude the possibility that PpcA can be replaced by one of its homologues to transport electrons from the inner to the outer membrane [17]. NMR direct electron transfer experiments suggest an equilibrium between ImcH and PpcA, explained by the overlap of the proteins’ redox potentials. The reduction potential values of ImcH hemes are lower than -100 mV (supplementary information table 1), and in a range close to those of its physiologic partner. The other members of the NapC/NirT family have at least one heme with a higher reduction potential, which allows the oxidation of the MKH_2_ pool to be more favorable. In *Escherichia coli*, TorC high potential heme was shown to be essential for electron transfer to TorA [43]. CymA does not have a higher potential heme, however, electron transfer to periplasmic enzymatic partners such as FccA, NapAB, NrfA or ArrAB becomes favorable due to the thermodynamics of the reaction (supplementary information table 1). In *G. sulfurreducens* ImcH MKH_2_ oxidation and its electron transfer to the periplasm is likely to be driven by the high redox potential of the Fe oxides (higher than - 100 mV). From the periplasm, multiheme cytochrome porins and nanowires at the outer membrane level will ensure a rapid electron transfer from the periplasm to the external acceptor [44,45].

In cellular respiration, the free energy from redox reactions is converted into an electrochemical ion gradient across the inner membrane (H^+^/Na^+^), which is used to drive ATP synthesis through ATP synthases. In *G. sulfurreducens*, these electrons are transferred outside, and when high potential electron acceptors are present ImcH is essential for EET [11,13]. Our electrochemical results show that ImcH is electroneutral, as proposed for other members of the NapC/NirT family, since electrons and protons are released on the same side of the membrane, so its main role is to recycle the MK-pool [46]. However, ImcH contributes to the maintenance of a *pmf* since it releases protons on the P-side of the membrane.

The comparison of the ImcH structural model with NrfH allows the identification of a possible MKH_2_ binding site on the P-side of the membrane, close to heme 1, where conserved protonatable residues that can be relevant for MKH_2_ oxidation were identified, which may allow proton transfer to the periplasm. The remaining hemes transfer the electrons from the MKH_2_ pool to the solvent exposed heme 7, which is the likely site for interaction with physiological partner(s).

Acetate oxidation is common to all *Geobacter* species, and a redox loop composed by NADH dehydrogenase (Complex I) and ImcH (Figure 5D) can be a widespread feature in many exoelectrogenic bacteria. Here, the electrons from carbon metabolism are shuttled through NADH, which is oxidized at Complex I on the N-side of the membrane with proton pumping. On the P-side, ImcH oxidizes the MKH_2_ pool with H^+^ release to the periplasm, where it reduces PpcA to start EET. Other electroactive bacteria, including *G. sulfurreducens*, can use H_2_ and formate as energy source, and electrons are transferred to the MK-pool through membrane bound hydrogenases and formate dehydrogenases, respectively. Curiously, in *G. sulfurreducens*, the annotated membrane bound hydrogenases (GSU0782-0785) and formate dehydrogenases (GSU0777-0779) have an integral membrane subunit with ten TMH, without co-factors, homologous to the NrfD family, rather than a cytochrome *b* subunit. NrfD subunits can be associated with electrogenic, electroneutral or energy driven mechanisms [47]. An electrogenic mechanism can be envisaged for H_2_ or formate oxidation since the thermodynamics of the reactions allows it, but experimental evidence is still required.

Regarding the other inner membrane proteins from *G. sulfurreducens*, CbcL and CbcBA, they both have an integral membrane domain/subunit homologous to the di-heme cytochrome *b* family that may be involved in MKH_2_ oxidation. Their contribution to energy conservation is still unknown and will be highly dependent on which side of the membrane the MKH_2_ oxidation takes place. It is more likely that MKH_2_ oxidation occurs on the N-side of the membrane, with electrons being transferred to the P-side by the *b*-type hemes according to the membrane potential, with protons released on the N-side, favoring the electron transfer to lower potential acceptors on the periplasm, without contributing to the *pmf*. This would make ImcH the only respiratory membrane protein in *Geobacter* known so far that contributes to the maintenance of the *pmf*, as it was previously predicted when a higher cell yield was observed in strains lacking CbcL and CbcBA *(ΔcbcL*, *ΔcbcBA*) [48].

In conclusion, this study presents the spectroscopic and electrochemical characterization of the quinone-interacting cytochrome ImcH, its interaction with periplasmic cytochromes and provides evidence for how energy is conserved in *Geobacter* species when EET is performed to high redox potential terminal electron acceptors (or electrodes). By transferring electrons from the MKH_2_ pool to PpcA via ImcH, protons are released on the same side of the membrane, without dissipating the membrane potential and contributing to ATP synthesis. This work provides important insights into *Geobacter* respiratory proteins that can be modified in order to tailor their metabolism to improve the use of these organisms for biotechnological processes.

## Methods

### Cloning Geobacter sulfurreducens ImcH

The *imcH* gene from *G. sulfurreducens* (GSU3259) was cloned into the pBAD202/D-TOPO vector (Invitrogen) as previously described [49] using the primers listed in Table 1. This construct inserts a 30-aminoacid linker with a 6x His-tag coding sequence at the C-terminal (KGELKLEGKPIPNPLLGLDSTRTGHHHHHH) that facilitates protein purification. Cloning was performed according to manufacturer (Invitrogen) specifications, and the final vector (pBAD-Gs-ImcH) was inserted into *S. oneidensis* MR-1 by electroporation [50] and plated in solid Luria Broth (LB) supplemented with kanamycin (30μg/ml) medium.

**Table 1.**
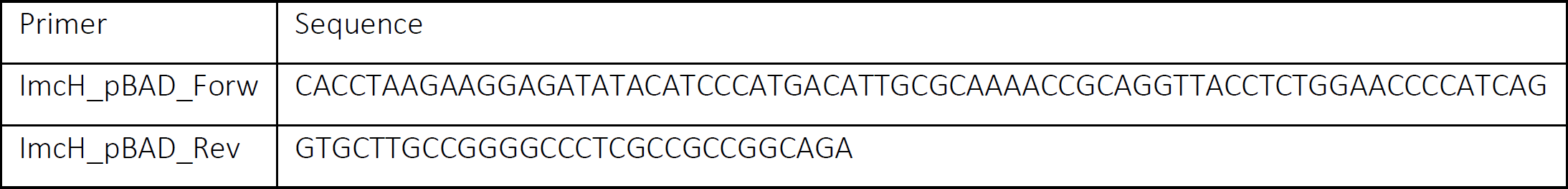
Primers used to clone ImcH from *G. sulfurreducens*.

### Protein production and purification

A single colony of *S. oneidensis* MR-1 carrying the pBAD-Gs-ImcH vector was inoculated in liquid LB selective medium and grown at 30 °C under constant agitation (180 rpm) until the culture reached a high OD _600 nm_ of 1.8-2.0. The best expression conditions were achieved by inducing gene expression with 5 mM arabinose and by lowering the temperature to 16 °C after induction, allowing the cells to grow overnight at 180 rpm. Cells were harvested by centrifugation at 11 300 *x g* at 8 °C for 10 minutes, ressuspended in 50 mM Tris-HCl pH 8.0 and disrupted using a French Press (10 000 Psi, three times). The membranes were collected through ultracentrifugation at 210 000 *x g* at 4 °C for 90 minutes, and extracted with n-dodecyl-β-D-maltoside (DDM). Briefly, the membrane fraction was incubated in 50 mM Tris-HCl pH 8.0 buffer containing 2% (w/v) of DDM stirring overnight at 4 °C, and the detergent solubilized proteins were then separated by ultracentrifugation at 210 000 *xg* at 4 °C for 90 minutes. For ImcH purification, the solubilized fraction was loaded on a DEAE Biogel (BioRad) column (Æ 2.6 ×;18 cm) equilibrated with 50 mM Tris–HCl pH 8.0 with 0.1% (w/v) DDM. The cytochrome rich fraction was eluted before increasing the ionic strength. This fraction was concentrated by ultrafiltration and loaded on an IMAC column (Æ 2.6 ×;14 cm) equilibrated with 50 mM Tris–HCl pH 8.0, 300 mM NaCl with 0.1% (w/v) DDM. A stepwise gradient of increasing imidazole concentration up to 500 mM was applied to elute the recombinant protein. Fractions were gathered according to the ratio absorbance between 410 nm and 280 nm (Asb 410/Abs 280). To obtain pure protein samples, the fractions containing ImcH were loaded on a Sephacryl S300 column (Æ 1.6 ×;90 cm) equilibrated with 50 mM HEPES pH 7.6, 150 mM K_2_SO_4_, 0.1 % DDM. Pure ImcH fractions were concentrated and frozen at −80 °C prior to use. For all chromatographic steps, protein purity was evaluated through UV-visible spectra and SDS-PAGE stained for total protein and heme staining [51].

*G. sulfurreducens* periplasmic cytochromes: PpcA, PpcB, PpcD, PpcE and PccH were heterologously produced in *Escherichia coli*, as described previously [31,34,52,53].

### Biochemical analysis

Total protein concentration was determined with the BCA colorimetric assay, and heme iron quantification was performed by the pyridine hemochrome method [54]. N-terminal sequence was performed by Edman’s degradation protocol using in house facilities. A molar extinction coefficient for ImcH was determined for the reduced state (*ε*_552nm_=148 mM^−1^cm^−1^) based on protein concentration. The ImcH oligomeric state was determined by size exclusion chromatography on a Sephacryl S300 (Æ 1.6 ×;90 cm) column equilibrated with 50 mM HEPES pH 7.6, 150 mM K_2_SO_4_ with 0.1 % (w/v) DDM using ferritin, glucose oxidase, albumin and chemotrypsinogen A as standards, mixed with blue dextran to determine the column dead volume.

#### Analytical centrifugation experiments

Sedimentation velocity experiments were performed using a Beckman Optima XLA-I analytical ultracentrifuge equipped with scanning absorbance optics. Measurements were performed at 20 °C and 40 000 rpm with 1.3 µM ImcH solubilized in 10 mM HEPES pH 7.8, 100 mM KCl, 0.1 % (w/V) DDM or 10 mM HEPES pH 7.8, 10 mM KCl, 0.1 % (w/V) Triton X-100 with 0, 20, 40 and 50 % (V/V) D2O. Each buffer density and viscosity was calculated with SEDNTERP [55]. Absorbance was recorded at 410 nm every 2 min. Samples were removed, migrated in SDS-PAGE gels that were stained for heme showed no evidence for PioA degradation. Data were fitted using the *c*(s) distribution analysis in the software SEDFIT [56]. The molecular weight*s* ImcH species within the c(s) distribution were obtained by fitting the data to Svedberg equation [55] and assuming a constant frictional coefficient

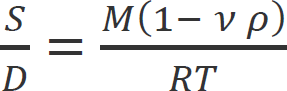

where *S* is the ImcH sedimentation coefficient; *D* the ImcH diffusion coefficient; ν the ImcH partial specific volume, *ρ* the solvent buffer density, *R* the gas constant and *T* the temperature.

Spectroscopic measurements

#### Kinetic experiments using stopped-flow

Kinetic experiments were achieved using a stopped-flow equipment (KinetAsyst SF-61 DX2, TgK Scientific) installed inside an anaerobic glove box with a N_2_ atmosphere. The temperature of the experiments was kept at 25 °C using an external circulating bath and changes in the absorption spectra were monitored by a diode array in the wavelength range 350–700 nm. Reduction experiments with menadiol were performed by fast mixture of a 2.6 μM ImcH sample equilibrated in 50 mM HEPES pH 7.6, 150 mM K_2_SO_4_ with 0.1 % (w/V) DDM with 100x molar excess of menadiol solution prepared in the same buffer. Menadiol was prepared by dissolving menadione in a concentrated ethanolic solution, reducing it with metallic zinc and quantified spectroscopically *ε*_(245nm–260nm)_ = 44.6 mM^−1^cm^−1^ [57]. Data were analyzed with Kinetic Studio software.

#### Potentiometric titrations of ImcH

ImcH potentiometric titrations were performed in an anaerobic glove box (98 % N_2_, 2% H_2_) at 20 °C in 50 mM HEPES pH 7.6, 150 mM K_2_SO_4_, 0.1% DDM. The solution potentials were measured using a combined Pt/Ag/AgCl electrode, calibrated with quinhydrone-saturated solutions at pH 4 and 7. A 3.8 μM ImcH solution was prepared with redox mediators: potassium ferrocyanate, 1,2 naphtoquinone-4-sulphonic acid, 1,2 naphtoquinone, 1,4 naphtoquinone, duroquinone, indigo tetrasulfonate, phenazine methasulfate, phenazine, 2,5-hydroxy-p-benzoquinone, 2-hydroxy-1,4 naphtoquinone, safrinine, benzylviologen and methyl viologen, all in a final concentration of 0.8 μM. Sample reduction and oxidation were achieved with the addition of small volumes of concentrated sodium dithionite or potassium ferrocyanide solutions, respectively, prepared in the same buffer. After each addition, the redox potential was allowed to equilibrate and a UV-visible spectra was recorded. ImcH reduced fraction was determined using the following equation:

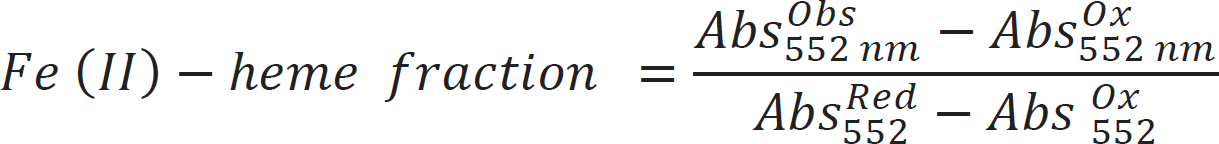

where Abs^Ox^ and Abs^Red^ are the absorbance at 552 nm for fully oxidized and fully reduced protein, respectively, and A^obs^ is the measured absorbance.

#### EPR spectra of ImcH

EPR spectra were recorded on a Bruker EMX spectrometer equipped with an ESR-900 continuous flow helium cryostat at 8 K under the following conditions: microwave frequency 9.39 GHz; microwave power 2 mW; modulation amplitude 1 mT; gain 6.32 × 10^4^. Two ImcH samples solubilized in 50 mM HEPES pH 7.6, 150 mM K_2_SO_4_ and 0.1% (w/V) DDM at the concentration of 118 μM were analyzed, one as purified and another reduced with excess of sodium dithionite (30x molar excess).

#### NMR experiments of ImcH

NMR experiments were performed on a Bruker Avance III 600 MHz spectrometer at 25 °C. Spectra were acquired with 2k increments for the protein-protein interaction in the oxidized state, 1k increments for the electron transfer experiments and with a sweep width of 70 kHz. All spectra were processed using TopSpin (Bruker). ImcH samples were buffer exchanged to 50 mM potassium phosphate buffer pH 7 in D_2_O with 0.1 % DDM. PpcA samples were lyophilized and resuspended in the same buffer. The DDM concentration was determined by 1D ^1^H-NMR and adjusted to maintain the experimental conditions. For protein-protein interaction studies, a 50 μM PpcA sample in 50 mM potassium phosphate buffer pH 7 with 275 mM DDM was titrated with ImcH at 1:1 and 1:2 ratios and the chemical sift changes of the heme methyl resonances were monitored. To assess the ImcH-PpcA electron transfer reaction, a 50 μM ImcH sample containing 150 mM DDM was reduced with a catalytic amount of hydrogenase in the presence of H_2_ inside an NMR tube sealed with a gas tight cap. H_2_ was flushed out with argon before the addition of increasing amounts of a PpcA concentrated sample in the molar ratio of 1:1, 1:2 and 1:3, inside a glovebox under N_2_ atmosphere. A 1D ^1^H NMR spectrum was acquired after each addition.

### Electrochemical experiments

All electrochemical measurements were performed inside an anaerobic glove box (N_2_ atmosphere), at room temperature (23 °C) using an electrochemical cell in a three electrode system: a working electrode (glassy carbon or pyrolytic graphite), a platinum wire as a counter electrode and a saturated calomel electrode (SCE) as the reference. Electrodes were connected to a PGSTAT204 potentiostat (Autolab, country) with data acquired by the manufacturer’s software (NOVA). Working electrodes (∅ = 0.3 cm) were immersed in a diluted nitric acid solution, rinsed in deionized water, hand polished in different grain size alumina (1, 0.3, 0.05 μm). Catechol was grafted to the glassy carbon working electrode as described in Lebègue *et al.* [58]. Briefly, an electrolyte solution containing 1 mM 4-nitrocatechol and 3 mM NaNO_2_ prepared in an aqueous acidic solution (0.1 M HCl) was degassed under argon and used in two cyclic voltammograms, between -900 and + 100 mV (vs SCE), at a scan rate of 50 mV.s^−1^. Afterwards, several cyclic voltammograms were acquired in the previous conditions using 1 mM HEPES pH 7.6, 150 mM K_2_SO_4_ buffer as electrolyte solution to observe the grafted catechol redox signal. The electrolyte buffer solution was replaced to discard desorbed catechol molecules and the electrode was poised at -900 mV (SCE) for 90 seconds prior to any experiment. Differential pulsed voltammetry was performed between -900 and +700 mV (vs SCE) with a 5 mV step, 200 mV modulation amplitude, 0.1 seconds modulation time and 1 second interval time (approximately 5 mV.s^−1^). ImcH was dissolved in a lipid solution containing phosphatidylcholine (PC) and menaquinone MK-4 to a final concentration of 3.9 mM PC 1 mM MK-4, 14 µM ImcH. This solution was incubated overnight at 4 °C with 500 mg of biobeads to remove the detergent. The catechol grafted electrode was immersed in the ImcH-MK-4-Lipid solution and immobilization was made through the solvent casting technique. An equivalent solution without ImcH was used for the control experiment.

To evaluate electron transfer between ImcH and PpcA, electrochemical experiments were performed by immobilizing ImcH (4 μL of a 35 µM concentrated sample) to a freshly polished pyrolytic graphite electrode through the solvent casting technique. The electrode was then immersed in a 250 mM potassium phosphate buffer solution at pH 7.0 and cyclic voltammograms were recorded between -900 and 100 mV (vs SCE) at different scan rates, from 10 to 100 mV.s^−1^. PpcA was added to the electrolyte solution in a final concentration of 1.5 µM from a concentrated sample equilibrated in the electrolyte buffer. Control experiments were performed using the bare electrode and PpcA in solution. Voltammogram subtraction was made by the direct subtraction of the second scanned voltammograms obtained at the same scan rate. Potential values are presented vs SHE by addition of 241 mV.

### Surface plasmon resonance experiments

Surface Plasmon Resonance experiments were performed on a BIAcore 2000 instrument (GE Healthcare). All assays were performed at 25°C. Before each experiment, the NTA sensor chip (Cytiva) was preloaded with Ni^2+^ through the injection of a 500 µM NiCl_2_ solution. The assays were performed with 10 mM HEPES (pH 7.4), 150 mM NaCl, 50 μM ethylenediamine tetraacetic acid (EDTA), and 0.01% (w/v) DDM running buffer. Pure ImcH (50 nM) was bound to the NTA sensor, by the C-terminus 6x His-tag to an activated flow cell at a 5 μL/min flow rate to have an immobilization of around 150 resonance units (RUs). Another activated flow cell was similarly treated with running buffer in the absence of ImcH (control cell). Interaction experiments with purified *Geobacter* periplasmic cytochromes PpcA, PpcB, PpcD, PpcE and PccH was performed with a 180 s injection at the concentration 500 nM at a flow rate of 15 μL/min, followed by a 400 s dissociation period. Interaction experiments with different amounts of PpcA were performed with a 360 s injection of increasing concentrations of PpcA (range from 150 nM to 2000 nM) at a flow rate of 15 μL/min, followed by a 400 s dissociation period. Before a new cycle of surface activation and immobilization, the NTA-chip was regenerated with a 350 mM EDTA in running buffer solution for 1 min at a flow rate of 10 μL/min. Sensorgrams were obtained by subtracting the unspecific binding to the control flow cell in order to remove buffer artifacts and normalizing to the baseline injection. Each experiment was performed in duplicate. Equilibrium dissociation constant K^D^ was determined from duplicate experiments and according to the following equation 2:

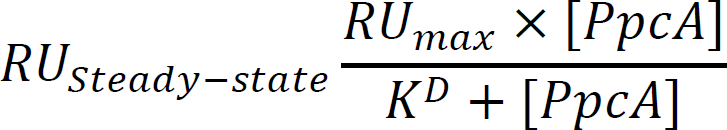

### ImcH AlphaFold model

*G. sulfurreducens* ImcH homology model was generated with AlphaFold2 [59]. The initial heme placement was manually optimized with Crystallographic object-oriented toolkit-Coot [60]. ImcH *c*-type hemes were fitted in Pymol using a self-generated library of *c*-type hemes and their corresponding Cys for the covalently binding of the heme vinyl groups, and His for the heme Fe coordination. The Rosetta software was used to optimize sidechain packing and carry out all atom energy minimization of the model [61]. This process was restrained using manually generated constraints for interatomic distances and bond angles for heme bound Cys and both proximal and distal iron atom ligands. Hemes were numbered according to their appearance in the protein primary sequence.

## Acknowledgments

This work was financially supported by Fundação para a Ciência e Tecnologia (FCT, Portugal) through grant PTDC/BIA-BQM/29118/2017 2017 (AGD), EXPL/BIA-BQM/0770/2021 (LM) and PTDC/BIA-BQM/4967/2020 (CAS), R&D units MOSTMICRO-ITQB (UIDB/04612/2020 and UIDP/04612/2020) and UCIBIO (UIDP/04378/2020 and UIDB/04378/2020), and associated laboratories LS4FUTURE (LA/P/0087/2020) and i4HB (LA/P/0140/2020).

