## Supplementary information for "Role of the inner membrane cytochrome ImcH in *Geobacter* extracellular electron transfer and energy conservation"

Supplementary table S1 – Details of the NapC/NirT family.

| Inner membrane | | | Periplasm | | | | Reference |
|---|---|---|---|---|---|---|---|
| | Number hemes | Hemes' reduction potential (mV) | Physiologic partner | Co-factors reduction potential (mV) | Pathway | $E^{0'}$ (mV) | |
| ImcH | 7 | -358<br>-326<br>-274<br>-150 (x4) | PpcA | -171<br>-119<br>-60 | EET | > -100 | [1,2] |
| CymA | 4 | -265<br>-240<br>-190<br>-110 | STC | -243<br>-222<br>-189<br>-171 | EET | | [3,4] |
| | | | FccA | -238<br>-196<br>-146<br>-102<br>-152 (FAD) | Fumarate reduction | +33 | [5] |
| NrfH | 4 | -300 (x3)<br>>0 | NrfA | -480<br>-400<br>-80<br>-50<br>+150 | Nitrite reduction | +340 | [6] |
| NapC | 4 | -235<br>-207<br>-181<br>-56 | NapAB | -160 (FeS)<br>-15<br>+80 | Nitrate reduction | +420 | [7,8] |
| DorC | 5 | -276<br>-185<br>-184<br>-128<br>-34 | DorA | Mo<br>+26/+200<br>W<br>-194/134 | DMSO reduction | +160 | [9] |
| TorC | 5 | -177<br>-177<br>-98<br>-98<br>+120 | TorA | n.d. | TMAO reduction | +130 | [10,11] |

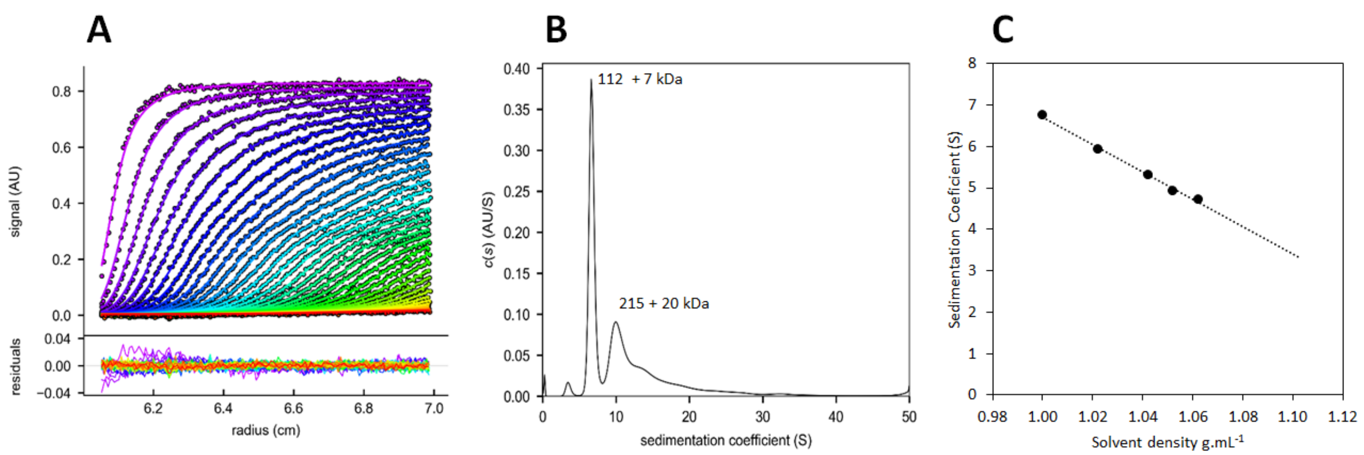


**Supplementary figure S1– Hydrodynamic analysis of ImcH solubilized in Triton X-100.** (A) Sedimentation velocity analysis of ImcH by monitoring the absorbance at 410 nm, (B) fit to the Lamm equation with residual absorption from the fitted data (lower panel from (A), (C) Extrapolation of the sedimentation coefficient (3.43 S) to the solvent density of 1.099 g $ml^{-1}$.

```
                                                  *       20         *       40         *       60         *       80         *      100
Geobacter sulfurreducens          ---------MTLRKTAGYLWNPISLIGFLLAVVATGLIIAFIAMEMITGI--DHPYIGLLVYFAFPGMLILGLILVPIGAWRVRNQRRTEVPE--EVPPY
Koribacter versatilis             -MPEDTGKIASDGKNKKLTRNVISLIGTTLAIVAAANILFLFLMDVMSPH--SSPYVGILAYMVMPAFLVLGLLMIPAGAWWERRRR--------GASSL
Ignavibacteria bacterium          ----------MKNKLPSSFYNPTSLAGAAIAAISFGLIVFLFLLELLGTK--QKPYMGIIAFVILPSILIGGVFLVIFGIYREHKREKHG-KQ--REKKF
Draconibacterium sediminis        --MKLPSS----------IRNWVSITGAVLAVFNLASILALFLLNVAFGF--GGQYIGLFIFIILPVFLIVGLLMIPLGMRIYRKKARLAEKEG-KRLNW
Opitutus terrae                   --MSTPSSPASPLKPRSAFNNWISAIGAVLSLGALFSFAFLVWMDFSQGE--KNPYLGILTYIVAPMFLMAGLALLFGGAWLQRRYAIKHAQ---NRPDR
Desulfuromonas acetexigens        MKRLSEIVGAFTRG---IARSRVSLIGAMIVTASAPFLFGAIFYDVLFHI--DNTYIAGAIYMLLGPAFIGGLILVFLGLFFFKGKEEVRLFTLDYLRDY
Geoalkalibacter subterraneus      MKKLTDIVSAFTKG---IARSRVSLVGAMVVTAVFPFLFGAILYDLTAHI--DNTYMAALIYMILGPAFIGGLVLVFLGLFFFKGKEEVRLFTMDYLRDF
Anaeromyxobacter dehalogenans     MP-SSRSGADWLR-------HPVSLAGAALATVSGVLVVVLFAASLLGLE--GGPYLGILAYLLLPGLLVAGLILVPVGAALERRRLVRLAARGAAEPPL
Geothrix fermentans               MS-LKERIRDLVRLCFQLGNNWMTLLGVALTTSSAFTLVWFWFMELTSPR-AVHPYVGILLFLALPALFLVGLVLIPVGILRTRR---KRRLAGESPTPL
Acidobacterium capsulatum         MGRLKQWEEEWLRPFLFYGNNWISLIGGAITTAAAFVLLGYWVVTLIGHGGSANPYTGIIFDFLLPALFVFGLVLIPIGMGWRRR---SLYRAGQIPSVY

                                                  *      120         *      140         *      160         *      180         *      200
Geobacter sulfurreducens          PRVDFNDPHKRRLFIFFVLASVIFVLIVSVASILGFEFTESTTFCGELCHVVMEPEHKAWQGSP-HARVKCVECHVGPGAEWYVKAKLSGLRQVWAVLTH
Koribacter versatilis             PTLDLNQPSQRSTVAFVVSFGIVFMMISAVGSYKAYEFTDSISFCGELCHTVMHPEFTAYQQSP-HARVACVECHVGSGASWYVKSKMSGLRQVYAATLN
Ignavibacteria bacterium          PTIDLNNPKHRAAFTFFSIGTILLLLFSAFGSYQAYEFTDSDEFCGEICHEVMEPEYTAYQFSP-HAKVGCVQCHIGSGAEWYVRSKFSGAYQVYSVLFN
Draconibacterium sediminis        PILDFNNLATRNASTIFIVGTIFLLIISSVGSYEAFHYTESVEFCGKLCHEVMEPEYVTYHGSS-HERVACVECHVGSGASWYVKSKLSGLYQVYSVLAN
Opitutus terrae                   WQVDFSNRRQRRIALGFGFGGILFLVLSAFGSYQTYHYSESTQFCGQVCHEAMNPEFVTYQRGE-HARVDCVECHIGSGAEWFIKAKINGTHQLIAYALD
Desulfuromonas acetexigens        FTDPHKFNRMRKLVFFAVFLTGINILIVSLLGYRTYHYMESVAFCGEFCHTVMNPERTAYLNSP-HSRVTCVECHIGAGADWFVKSKISGARQLFAVALN
Geoalkalibacter subterraneus      FTDPTKFNKMRKLVFFAVFLTGVNIFIMGLLAYRGYHYMESNGFCGQFCHTVMNPEYTAYQNSP-HSRVNCVECHIGSGATWFVKSKISGARQLAAVAFG
Anaeromyxobacter dehalogenans     PVMDLNHPRTRRRLLVFLGLTTVNLLLLGIASYKGLEVMDSPAFCG-SCHSVMDPEASAHRRSA-HARVACVECHIGPGASWFVKSKLSGAWQVVSVALD
Geothrix fermentans               QKVDFTQPGIRRLLLVVGALTFVNVGILGTAGLKGVEYMDSNRFCGLTCHTVMSPEYTAFLNSP-HSRVGCAQCHIGHGAPALVRAKISGTRQLFAVAFG
Acidobacterium capsulatum         PQVDFRDPKFRHAVDFVVVATFINFVIVGTASYRGVAYMDQASFCGTSCH-VMQPEWVAYHHSQFHTNVACVECHVAPGIPGYIHAKTNGTKQLLMVLFH

                                                  *      220         *      240         *      260         *      280         *      300
Geobacter sulfurreducens          SYHFPIATPIENLRPARDTCEQCHWPEKFYSGRQRVFYHYAPNKENTPREINMLIKIGG-TPKSPHAMGIHWH--IGTEVTYIARDRKRLDIPYVAVKQK
Koribacter versatilis             TFPRPIPSPVANLRPAAQTCEQCHWPKKFWGAQLKTIYHYGYDEKNTPRVVNLLVKTGGGDANNGTAMGIHWHMNIANKISYIS-DERRENISYVKAVNQ
Ignavibacteria bacterium          KYSKPIPTPIENLRPAQETCEQCHWPKHFFNEKQLVNTYYLSDENNTEWTIDLLIKIGGGNVEAGPTSGIHWHMNIAHEVNYVTLDSLRMIIPWVQSKSA
Draconibacterium sediminis        KYPQPIPTPIANLRPARETCEECHWPEKFYDNKMRVKHSFLTDEENTEHIVHLQVKTSTQVTPQGVIKGIHQHISPEVKIEYKALDEKRQIIPWVKYTNT
Opitutus terrae                   NYNRPIATPLHNLRPAQDICEKCHWPEKFHGNIEINFDHFLSNKKNTPYTARMLMHVN-ASRPGGPAGGIHWHVNENEKVEYYAADAKRQEIPWMRVTNV
Desulfuromonas acetexigens        TYPRPIATPVHGLRPARDTCEECHRPEMFHGDKLVIKDKFLEDEQNTNVKTVLLMKVGSAGDRTSSSHGIHWHVAAENKIFYRPSDHSRMKIPEVIQAAE
Geoalkalibacter subterraneus      TYPRPIETPVHGLRPARDTCEECHRPEKFHGDKLVIRDKFLDDEENTHVQSVLLMKIGSAGDRTTSAHGIHWHVAPENKIIYKAADWQRTVIPEVVLQQA
Anaeromyxobacter dehalogenans     LYPRPIPTPVQNLRPARDTCEQCHWPTKLVGDRLKVITRHAEDAGSTPLRTVLVVHVGGAQGLGARTRGIHWHVDPGVRIRYLADEKR-ETIGTVELTGP
Geothrix fermentans               TYSRPIPSPVEHLRPARETCEQCHWPQKFTDEKLLIRTKYADDETNTPATSILLLKIGGHTPNGT-TGIHGRHLDAGERISYISTDHRRQEIPKVTYRDD
Acidobacterium capsulatum         SYPRPIMA-DNKLPPPTVTCERCHSPTRYAGDQLVVKNTFGDDEKNTLTHTVLVMHVGGRNAAGVYSGIHGSHYK--EKIEYIATDASDQTIPWVAVIHP
```

```
                                       *       320         *       340         *       360         *       380         *       400
Geobacter sulfurreducens      D-GSIVEYMDTEKPLTREEIAK----------------------------------------------------------AEKRRMDCIDCHNRPTHIYR
Koribacter versatilis         K-GEVTEYFPKDNPPSADMVAN----------------------------------------------------------TAQRNMDCIDCHNRPSHIYV
Ignavibacteria bacterium      D-GKVTIYRNTELNISDSQVEK----------------------------------------------------------LEKRRMDCIDCHNRPSHIYH
Draconibacterium sediminis    KTGEEYIYTDEMSMVSEAELDS----------------------------------------------------------LETRVMDCLDCHNRPSHNYN
Opitutus terrae               KTGASRIYRTD-EFTGEPPVD------------------------------------------------------------QIRVMDCMDCHNRPAHMFP
Desulfuromonas acetexigens    DGSLLVYRSSEANAELQASGKAGLQSLRSEQVAVADSGQSHPASRAASGRWIAGPKGKQSERPPEVKDIRAYAEQGGYVMEDLREMDCIDCHNRPTHIYL
Geoalkalibacter subterraneus  DGTEKVFR-TRTDSEIEH--------------------------------------------------------------ADVREMDCIDCHNRPTHIYL
Anaeromyxobacter dehalogenans DG-----ARATFRSKAPAG------------------------------------------------------------EARWREMDCVDCHNRPTHVYR
Geothrix fermentans           NGQLVEYVTEDFKKLSPEQL----------------------------------------------------------AKATERKMDCIDCHNRPTHTFE
Acidobacterium capsulatum     DGTKTVYAVAGFHPAANQ---------------------------------------------------------------KPRVMGCIDCHNRAAHSFA

                                       *       420         *       440         *       460         *       480         *       500
Geobacter sulfurreducens      SPAREMDEHIVSG-QIDAGLPYIKKVAVEILEQPYKSKEEAHAAIEAKLPEYYAKNFPEVAKVKAAAINQAVAHVKDIYSRNFFPRMKVTWSTYPNHIGH
Koribacter versatilis         PPDRSVDNAMVAG-KIDPSLPFVKQQAVTILTKDYKTSAEALNTISVEFPKYFQTTYPQVAQQKAQQINDATAELQRIYKNSFFPEMKVSWKTHPNHIGH
Ignavibacteria bacterium      PPARSVNHLIKLG-WIDKRLPYIKSLSVHALENEYTSKQVALDSIKSTILDFYSANYPQVFIEKKTSINKAVEEVQKIYSRNYFPSMGVSWRKFPEHIGH
Draconibacterium sediminis    APQNFIDQSMAEG-KISTDLPAIKLAAMMALYQDYPTKDTAMIAIENQITDWFESGYPEVYETKKAEIDQAIAAIQEDYSNNIFPFMKANWKAYPNNIGH
Opitutus terrae               TANEAVERALLTG-TLSTALPNIKRDAVNAMTQEDITTS---AGAPEKIAAFLTKKY-----AGHPDLPGAIAAVQSLYATTIFPERKADWRVYPNNIGH
Desulfuromonas acetexigens    PPGVAVDNQIMSG-QIDVSIPYIKKKAMEVVLREYSSQDQAKETIAAEIREFYGTNYPQVS---AEALERSIAGIQQAYAENVFPEMNIQWGTYTNHLGH
Geoalkalibacter subterraneus  PPGRAIDNKILEG-KIPSEIPFIKKKAMEVVQKEYATQDEARTRIATDLTAWYKQNYPEIGRDNPEKLQKAITGIQEAYSENVFPEMKIEWNTYTNHIGH
Anaeromyxobacter dehalogenans SPEDEVDAAVAAGRIDAAALPFARREALKALRAGYPSHEAARAGIAAALGAFYAG--PEGGGAPREAVERAASELGDAWARNVWPSMRIAWGTYPSLLGH
Geothrix fermentans           LPDRAVDKSMAKG-LIATDLPYIKKKAVELLKTEYASQQEAGSKIVLGINEFYRTTYPDVFKHKRAQVDAAAETVKTIYLRNVFPEMKLTWGTHPNNIGH
Acidobacterium capsulatum     SPGDAMDRAMAGG-ALPASLPYLHKEGMALIQANYVSQGEAATKIPKALEAYYRTNYPAIWSGQSSEVKQAGQELVALYEQNVFPFMKVTWGTHPNNIGH

                                       *       520         *       540         *       560         *       580         *
Geobacter sulfurreducens      FYT----PGCFRCHDGKHKTSTGKIISK-DCNMCHEMIGQK--GENIPEGK-VVKEFVHPADIGDALYNVNCSDCHMAAAEDSAGGEGPGKH
Koribacter versatilis         FYY----NGCFRCHDGNHVSKSGKVISK-DCKICHTMLSQEENGVQISQST-SGVEFKHPVDIGD-LSQVNCSDCHTGGAQ-----------
Ignavibacteria bacterium      LYA----PGCFRCHDGKHVNEEGKVLSR-DCNVCHTILAQKFEKDELRLSL-GGIDYQHPVDVGNAWKEINCSECHEKK-------------
Draconibacterium sediminis    MES----DGCYRCHNDRHATTENKTISK-DCSLCHNIIAQGTPDSMQYSTAFEALDFQHPVDVAEAWRSEMCSMCHTALY------------
Opitutus terrae               KDW----PGCFRCHDDKHKTASGQTVSASDCTSCHTIIAQGKGEELALLSA-KGLEFEHPGGEYDAELT--CADCHNGGVQK----------
Desulfuromonas acetexigens    NDD----VGCFRCHDEDHETETGEVISM-DCDTCHTILAEDEADPEILRTLSGG--------------------------------------
Geoalkalibacter subterraneus  TEDFDFSIGCLRCHNDMHETESGEVISM-DCDTCHLVLAQDEEDPEVLRTLGGGY-------------------------------------
Anaeromyxobacter dehalogenans EAA----PGCFRCHDGDHETEDGRAIRA-DCDLCHEILAQEEKDPAVLKLLA----------------------------------------
Geothrix fermentans           EEA----VGCFRCHDGSHTAADGRTIAA-DCNTCHTILAQDEKDPKVLKDLGMK--------------------------------------
Acidobacterium capsulatum     MAS----PGCFRCHDGNHVTTGGTALTN-DCTVCHNIVAMGDPNPKPLAELGMTQ-------------------------------------
```

**Supplementary figure S2 – ImcH multiple sequence alignment**. Sequences were downloaded from the Joint Genome Institute (https://jgi.doe.gov/) and aligned with ClustalX 2.1. Conserved residues are colored in black except for the predicted heme-Fe axial ligands, histidines 137, 152, 161, 210, 256, 258, 316, 321, 423, 433, 453,474 and 491 (cyan), glutamine 179 (green) and conserved protonatable residues near heme 1 pocket Glu127, Lys173 and Tyr188 (red).

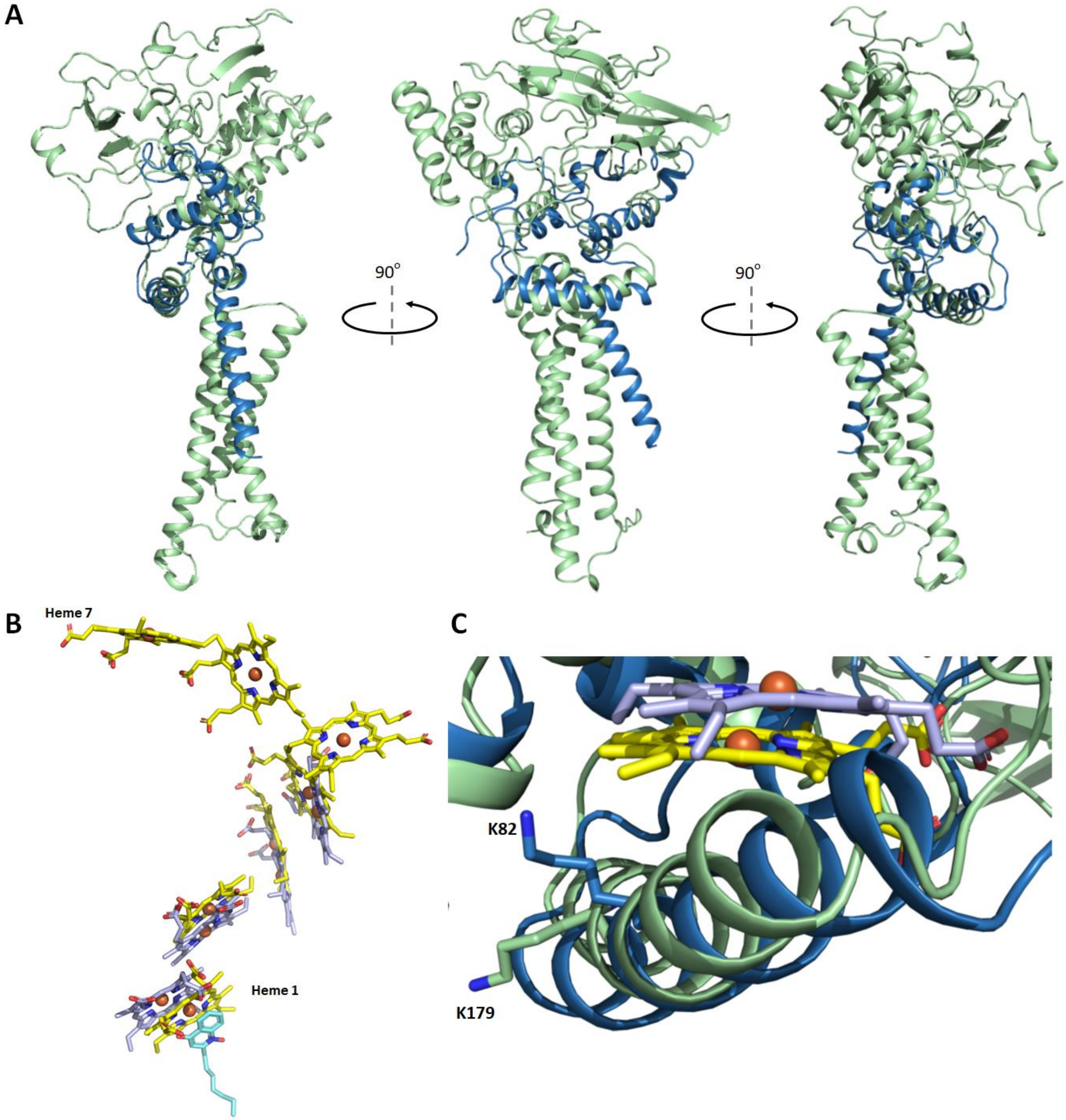


Supplementary figure S3 – *G. sulfurreducens* ImcH structural alignment with *D. vulgaris* NrfH. (A) The structural model of *G. sulfurreducens* ImcH (green) is superimposed on the structure of the NrfH subunit (blue) of the *D. vulgaris* Hildenborough cytochrome *c* nitrite reductase ($NrfA_4H_2$) PDB code:2J7 [12]. A cartoon representation is used to depict the backbone structure of the two proteins. (B) Superimposed hemes represented as sticks with carbons colored in yellow (ImcH), colored in light blue (NrfH), colored in cyan (HQNO), nitrogen in blue, oxygen in red, and Fe atoms as orange spheres. (C) Detail of the superimposed structures near heme 1 pocket, with lysine residues represented as sticks: K82 from NrfH (blue) and K179 from ImcH (green).

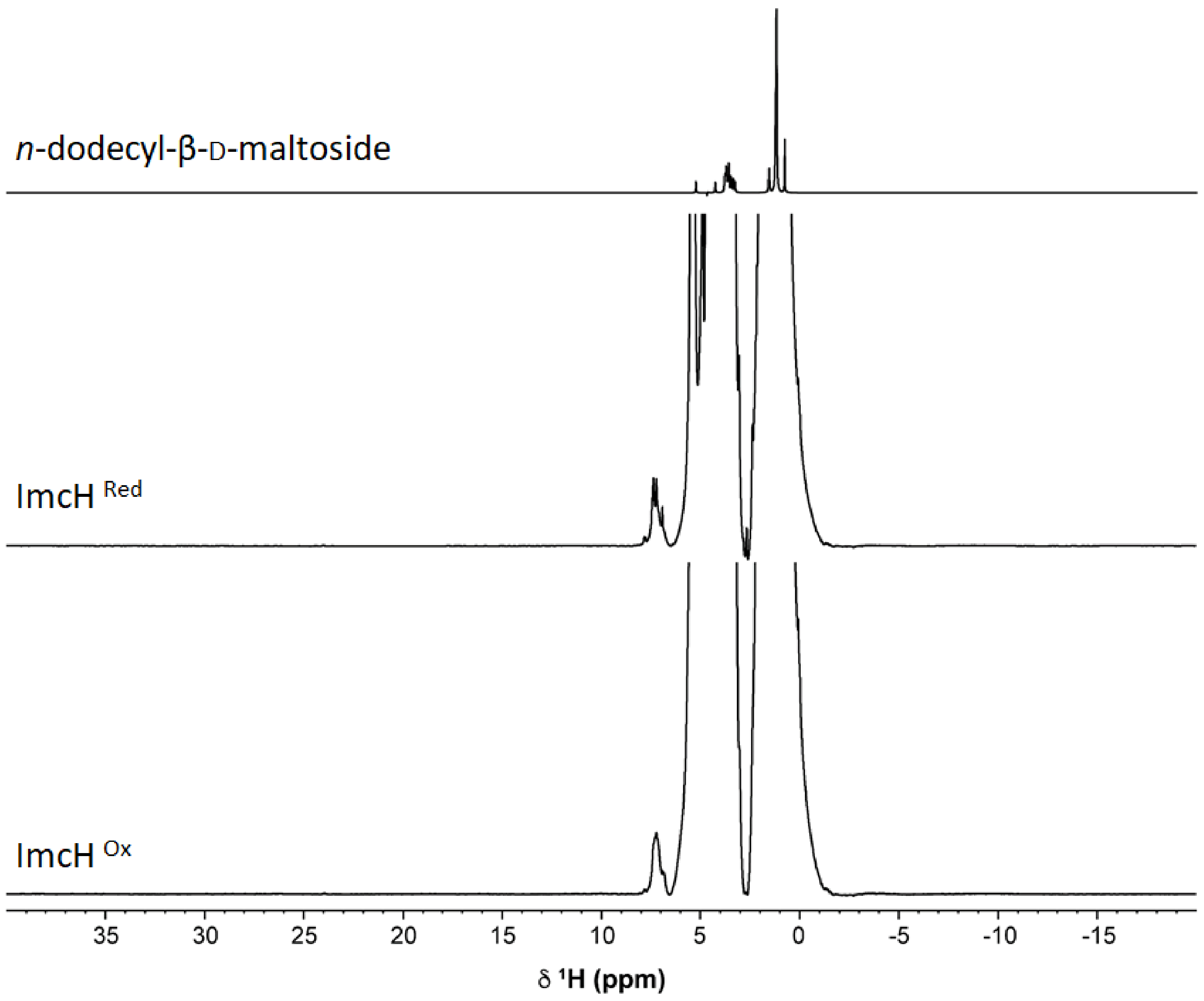


**Supplementary figure S4 – N-dodecyl-β-D maltoside and ImcH 1D $^{1}$H-NMR spectra.** 1D $^{1}$H NMR spectra of 100 mM DDM (top), and ImcH (50 µM) in the reduced and oxidized states (50 mM potassium phosphate buffer pH 7 and 150 mM DDM in $D_2O$). Both ImcH spectra are zoomed to reveal the protein signals.

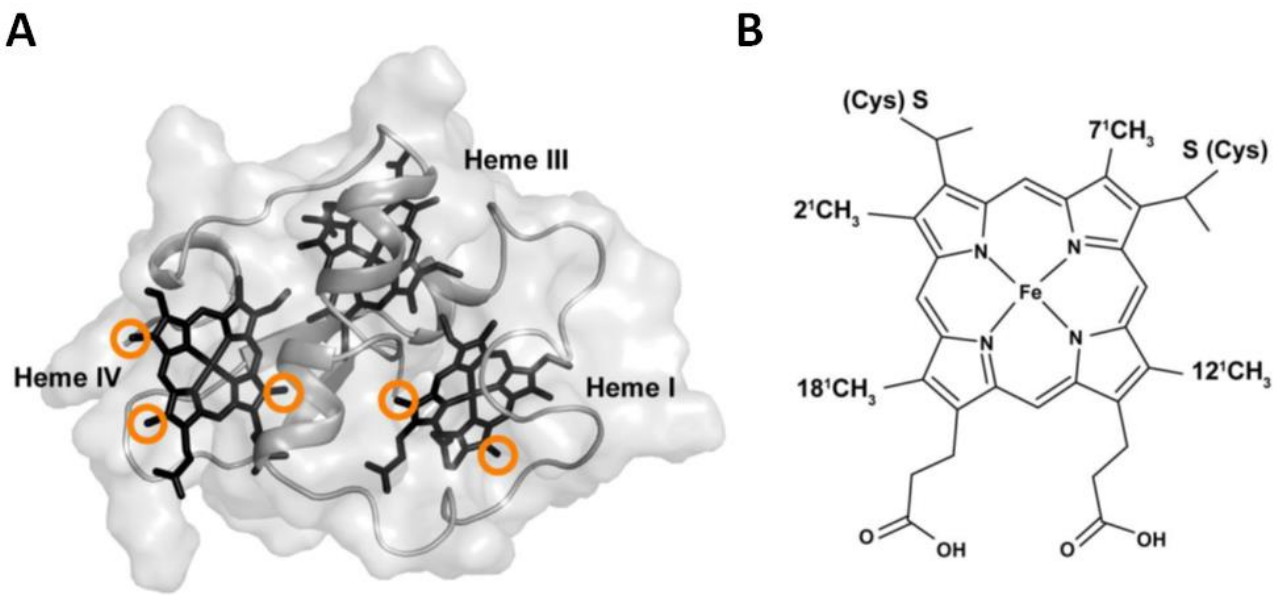


**Supplementary figure S5 – Structure of *G. sulfurreducens* triheme cytochrome PpcA.** PpcA structure (PDB 2MZ9) with hemes shown in sticks. Orange circles highlight methyl groups that show differences on the low-field 1D $^{1}$H NMR spectrum upon interaction with ImcH (see Figure 4C), (B) Schematic representation of the heme *c* with methyl nomenclature.

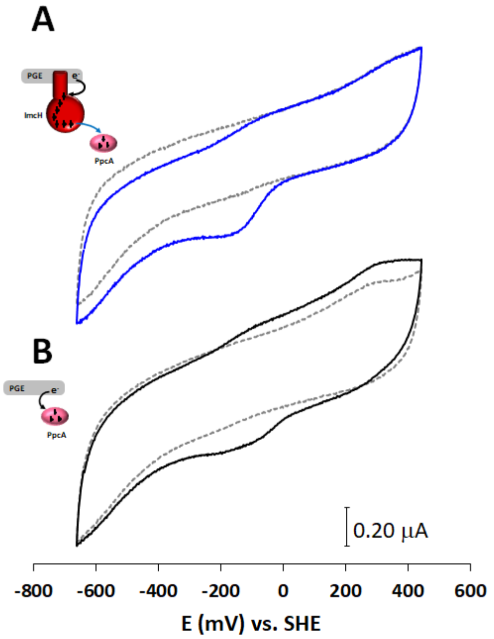


Supplementary figure S6 – Direct electron transfer between ImcH and PpcA monitored by cyclic voltammetry. Cyclic voltammograms performed at 100 mV s$^{-1}$ (A) with ImcH modified graphite electrode without PpcA in the electrolyte solution (grey dashed line) and with PpcA added to the electrolyte solution (blue line) (B) performed at 100 mV s$^{-}$ with bare graphite electrode without PpcA in the electrolyte solution (grey dashed line) and with PpcA added to the electrolyte solution (black line). Cyclic voltammograms of graphite bare electrode (grey line) and ImcH modified electrode (black line) performed at (C) 10 and (D) 100 mV s$^{-1}$.

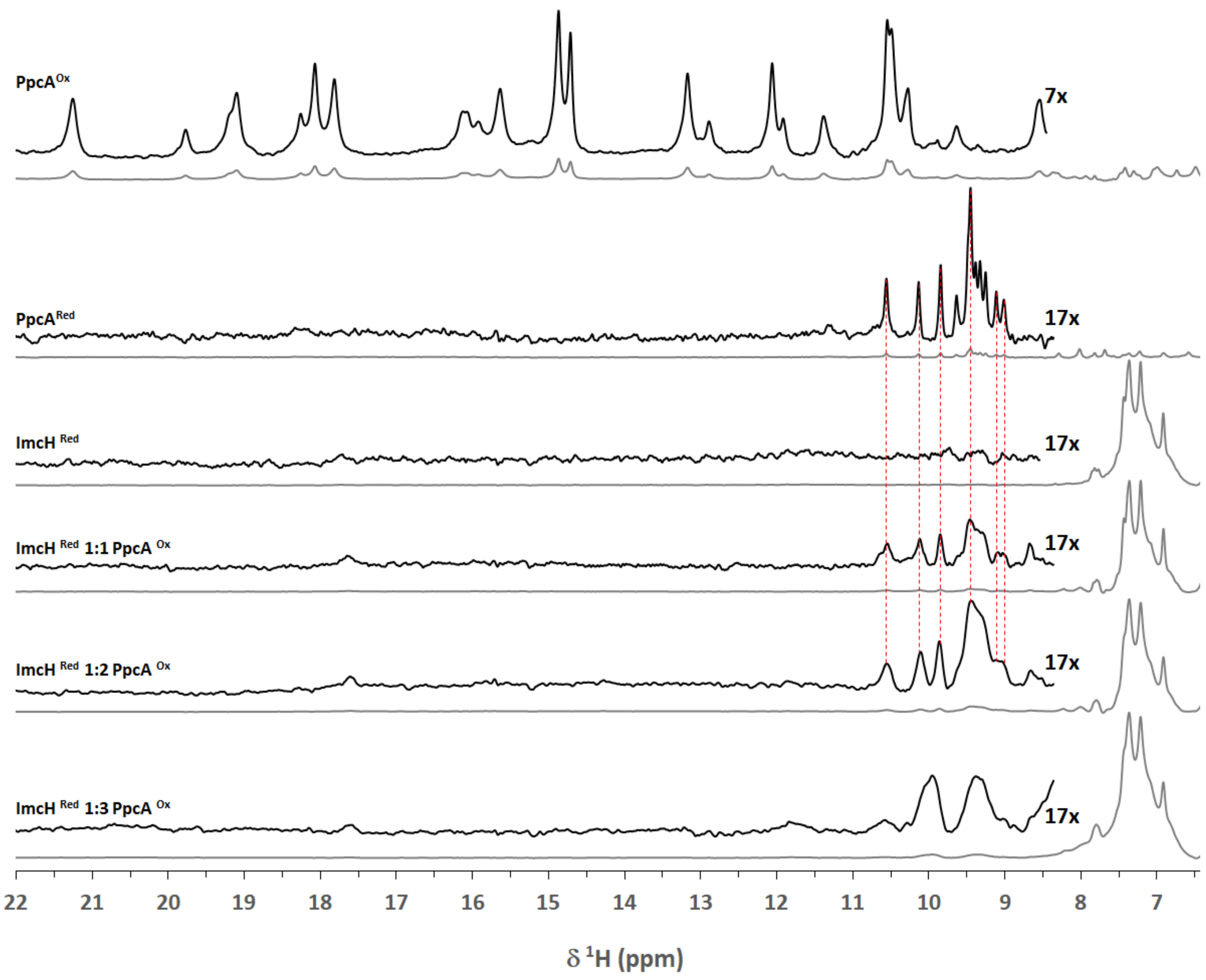


Supplementary figure S7 – Direct monitoring of the electron transfer reaction between ImcH and PpcA by NMR. Low-field 1D $^{1}H$ NMR spectra to detect the interaction between reduced ImcH and oxidized PpcA at increasing molar ratios. Spectra for fully reduced and oxidized PpcA, and fully reduced ImcH are displayed at the top. The vertical red dashed lines highlight the chemical shifts typical for reduced PpcA upon addition to reduced ImcH.
